# Untargeted Plasma Metabolomics Unveils Distinct Metabolite Profiles in Parkinson’s Disease Subtypes: A Focus on idiopathic REM Sleep Behavior Disorders

**DOI:** 10.1101/2024.05.02.592293

**Authors:** Sunjae Lee, Jihyun Kim, Jaewoo Baek, Ki-Young Jung, Yunjong Lee, Ara Koh, Han-Joon Kim

## Abstract

**Background:** Parkinson’s disease (PD) is characterized by diverse clinical presentations and etiological complexities, with rapid eye movement (REM) sleep behavior disorder (RBD) serving as a prodromal marker. While extensive unbiased metabolic profiling of plasma samples from PD subjects has been conducted to identify novel PD metabolic biomarkers, comprehensive metabolic profiling of PD subtypes based on RBD status remains limited.

**Methods:** We conducted a comprehensive metabolic profiling of PD subtypes at disease onset, considering the presence or absence of RBD, utilizing an untargeted metabolomics approach. Plasma samples were collected from subjects with PD with and without RBD at the initial stages of disease, idiopathic RBD, and healthy controls to elucidate similarities and differences among PD subtypes. Based on ordination analysis and metabolome-wide association study (Wilcoxon rank-sum tests and generalized fold changes), we identified specific groups of metabolites enriched in the PD_Only group and RBD groups (iRBD & PD_RBD+), with few metabolites shared between groups. Furthermore, pathway enrichment analysis (hypergeometric tests) identified specific groups enriched with metabolites from specific origins and associated biospecimens, as well as disease-associated metabolites. Finally, we evaluated the biomarker potential of the identified disease metabolites by ROC curves and proposed logistic regression models of key biomarkers and clinical parameters for predicting disease status.

**Results:** Metabolomic analysis revealed distinct metabolic profiles between PD subtypes with and without RBD. Our analysis confirmed previously reported PD metabolic markers, such as a reduction in caffeine and urate, as well as an increase in cortisol, secondary bile acids, and p-cresol sulfate. However, our stratified analyses based on the presence of RBD discriminated RBD-associated metabolites from those associated with PD_Only (without RBD). PD patients with RBD exhibited enrichment of gut microbial-origin metabolites, including secondary bile acids and p-cresol sulfate, compared to PD patients without RBD. Conversely, metabolites associated with neuro-psychiatric diseases were enriched in PD patients without RBD.

**Conclusions:** Our study elucidates the heterogeneous nature of PD subtypes, particularly differentiated with the presence of RBD. The metabolic features of PD with RBD subtype supports the “body-first” concept of PD pathogenesis originating from the gut.

## Introduction

Parkinson’s disease (PD) is the second most prevalent neurodegenerative disorder, characterized by abnormal α-synuclein aggregation, termed α-synucleinopathies. These pathologies, notably in dopaminergic neurons of the substantia nigra pars compacta, significantly contribute to the motor symptoms associated with PD due to the loss of dopaminergic neurons (1). PD patients also develop diverse spectrum of non-motor symptoms in the progression of the disease and even in the prodromal stages. The different origin and transmission sequence of a-synuclein pathology are supposed to be responsible for heterogeneous manifestation of diverse spectrum of PD symptoms. Indeed, accumulating evidence indicates that clinical and prodromal PD exhibit notable heterogeneity, prompting the identification of proposed subtypes (2).

Idiopathic REM sleep behavior disorder (iRBD), recognized as a prodromal manifestation of α-synucleinopathies, strongly correlates with PD development, but less than 50% of clinically manifest PD cases exhibit RBD at onset (2). Recently, based on neuropathological and clinical evidence, two subtypes of PD have been proposed: brain-first (initiating in the brain) and body-first (originating in the enteric nervous system, reaching the brain via the vagus nerve) (2–4). iRBD is linked to the body-first type, with initial α-synuclein aggregation in the gut affecting medulla and pontine structures before nigral dopamine neurons. In contrast, PD without RBD at onset is associated with the brain-first type, marked by the initial loss of nigrostriatal dopamine innervation but relatively intact sympathetic and parasympathetic innervation (2, 4). However, the robustness of these two subtype PD hypotheses requires further verification, and exploring alternative PD subtypes is critical for improved diagnosis and treatment based on distinct etiologies in this heterogeneous disorder.

A compelling association exists between gastrointestinal symptoms, notably constipation, and the onset of motor symptoms in PD, often manifesting many years prior (5). Several studies emphasize the intricate link between PD and the gut microbiota (6, 7). Building on the body-first PD subtype, a recent cross-sectional cohort study examined microbial signatures in healthy individuals, first-degree relatives of iRBD, iRBD, and PD with RBD (8). This study indicates that PD-like changes in gut microbial signatures commence in iRBD, supporting body-first PD subtypes (8). Given the potential different etiologies between PD with RBD at onset and PD without RBD at onset, further investigation is needed into microbial signatures between two groups of PD.

The gut microbiome can influence not only gut but also brain phenotypes through the production of bioactive metabolites, either directly or indirectly (9, 10). Among these microbial metabolites, short-chain fatty acids (SCFAs) have been demonstrated to be capable of promoting microglia activation and motor deficits, potentially through an indirect pathway (11). Extensive metabolomic profiling of blood samples from PD patients has been conducted to identify metabolic biomarkers for better characterization or early-stage diagnosis of PD (12–16). Some studies have provided insights into circulating blood metabolites of gut microbial origin, such as p-cresol sulfate and phenylacetylglutamine (14, 16). However, it is not clear whether PD subjects used in these studies represent mixed subtypes of PD (brain-first or body-first). To date, there have been no studies on untargeted metabolite profiling of PD subtypes in the early stages of disease.

In this study, we thus aim to dissect metabolic profiles of PD subtypes by performing untargeted metabolomics on neurologically healthy controls, iRBD, PD with RBD, and PD without RBD in the early stages of the disease.

## Methods

### Subjects

The subjects in this study were recruited from the Movement Disorders Clinic at Seoul National University Hospital (SNUH), comprising a total of 101 subjects (**Table 1**). Among them, 24 had PD without RBD, 25 had PD with RBD, 25 had iRBD without movement disorders or dementia, and 27 were controls (categorized as PD_Only, PD_RBD+, iRBD, and Control group, respectively). Patients with PD were followed for an average of 4.0 years after sampling to ensure the reliability of the diagnosis. iRBD were diagnosed using video-polysomnography in accordance with the third edition of the International Classification of Sleep Disorders. The diagnosis PD was established following the clinical diagnostic criteria for PD by the Movement Disorders Society (17). PD subjects were further categorized based on the presence or absence of RBD at disease onset, resulting in two groups assessed using RBD1Q (18): the PD_Only group and the PD_RBD+ group. The control group, free of PD or iRBD, was drawn from the SNUH biorepository, predominantly consisting of spouses sharing living environments with PD or iRBD patients. The study protocol was approved by the Institutional Review Board of the Seoul National University Hospital (IRB No. 2207-085-1340), and informed consent was waived by the board.

**Table 1.** Demographic summary of PD cohort in this study.

|  | **PD_Only** | **PD_RBD+** | **iRBD** | **Control** | ***P*** |
| --- | --- | --- | --- | --- | --- |
| **n** | 24 | 25 | 25 | 27 | - |
| **Men, n (%)** | 12 (50) | 13 (52) | 13 (52) | 19 (70) | 0.99 |
| **Age at sampling** | 62.8 ± 7.3 | 62.0 ± 5.7 | 63.5 ± 6.8 | 62.4 ± 7.1. | 0.90 |
| **HY stage at sampling** | 1.2 ± 0.4 | 1.4 ± 0.5 | NA | NA | 0.15 |
| **PD duration at sampling** | 1.5 ± 1.4 | 1.2 ± 0.8 | NA | NA | 0.70 |

Abbreviation: HY, Hoehn and Yahr; NA, not applicable; PD, Parkinson's disease. Data are presented as mean ± s.e.m or as proportions. *P* values were calculated using ANOVA for continuous variables and chi square test for categorical variables.

### Plasma collection and metabolite analysis

Blood samples were collected from the participants’ venous blood during visits. The samples underwent centrifugation, and the resulting plasma was stored at −80°C until analysis. Global metabolite profiling analysis on the plasma samples was conducted by Metabolon (Durham, NC) using ultra-high-performance liquid chromatography coupled to tandem mass spectrometry (UPLC-MS/MS). In summary, automated sample preparation was carried out using the MicroLab STAR® system (Hamilton Company). A recovery standard was introduced before the initial step in the extraction process for quality control. Proteins were precipitated with methanol under vigorous shaking for 2 minutes (Glen Mills GenoGrinder 2000), followed by centrifugation. The resulting extract was divided into five fractions: one for analysis by UPLC-MS/MS with positive ion mode electrospray ionization, one for analysis by UPLC-MS/MS with negative ion mode electrospray ionization, one for LC polar platform, one for analysis by GC-MS, and one sample was reserved for backup. Samples underwent a brief period on a TurboVap® (Zymark) to eliminate the organic solvent. For LC, the samples were stored overnight under nitrogen before preparation for analysis. For GC, each sample was vacuum-dried overnight before preparation for analysis.

### Statistical analysis of plasma metabolomics

Using untargeted metabolomics profiles after zero-imputations for missing values, number of detected metabolites were checked and compared by groups, such as control, iRBD, PD_Only and PD_RBD+ by Wilcoxon rank sum tests. Similarity of metabolomics samples were compared by non-metric multi-dimensional scaling (NMDS) by metaMDS function of R vegan package (version 2.6-4) and their associated loadings were also calculated by the envfit function of vegan library. Differentially abundant metabolite testing was performed by Wilcoxon rank sum tests, together with generalized fold change (R MicrobiotaProcess Package, version 1.11.5) (19, 20), an estimator of log-2 fold change with capturing the variation of posterior distribution of log-2 fold change. Visualizations of boxplots, scatter plots and bar plots were performed by R ggpubr package (version 0.6.0). Metabolites of given metabolomics profiles were annotated with Human Metabolome Database (HMDB) (21) and origins, associated biospecimen and disease metabolite information were obtained from HMDB (https://hmdb.ca/) (as of 6th Feb 2024). Overlaps of significantly differential metabolites with above-mentioned metabolite classes (origins of metabolites, associated biospecimen and associated diseases) were checked with hypergeometric tests. For PD_Only metabolites (POM) and RBD metabolites (RM), we checked BBB permeability by lightBBB prediction program (22). AUROC based on the sensitivity and specificity were calculated with R pROC package (version 1.18.4). logistic regression models using key metabolites, sex, and age were generated using glm function (family = binomial) of R stats package (version 4.2.1) from randomly assigned training and test datasets (ratio of data 2:1, respectively)

### *Clostridioides difficile* abundance and *hpdB/C* abundance analyses

Publicly available metagenomic data of both Parkinson’s disease patients (n=491) and neurologically healthy elderly controls (n=234) were obtained from BioProject ID PRJNA834801 (23). To ensure data quality, the Nextera XT adapter and PhiX genome contamination were removed using BBDuk with the following parameters: ‘ftm=5 tbo tpe qtrim=rl trimq=25 minlen=50 ref=adapters,phix’. Subsequent reads were aligned to the human reference genome (GRCh38.p14) using BBSplit with default parameters to eliminate human contamination. To estimate the presence of authentic p-cresol producers, the abundance of *Clostridioides difficile* was assessed. Quality-controlled sequences underwent taxonomic profiling using MetaPhlAn 4 (v4.0.6, mpa_vOct22) with default parameters (24). For profiling functional gene contents, a species-level metagenomic functional analysis was conducted using HUMAnN3 with default parameters (25). The resulting counts based on UniRef90 (v201901b) and ChocoPHLAn 3 database were converted into Kyoto Encyclopedia of Genes and Genomes (KEGG) orthologs. To quantify the abundance of genes encoding the p-cresol-producing enzyme, hpdB/C, the specific gene identifiers K18427 (4-hydroxyphenylacetate decarboxylase large subunit) and K18428 (small subunit) were targeted.

## Results

### Clinical characteristics and plasma metabolic features of the cohorts in this study

Clinical characteristics of subjects were shown in Table 1. A total of 101 plasma metabolomics samples were generated, comprising 24 PD without RBD at onset (PD_Only), 25 PD with RBD at onset (PD_RBD+), 25 iRBD, and 27 healthy controls (**Figure 1A**). All clinical groups showed no substantial differences in their demographic features, including age, gender, Hoehn and Yahr (HY) stage, and duration of disease at blood collection. Patients diagnosed with PD were in the early stage of disease, with a mean disease duration of 1.3±1.1 years from symptom onset. In the PD_Only and PD_RBD+ groups, 18 (75%) and 19 (76%) patients, respectively, were not taking anti-parkinson medications. Four patients in each group were taking levodopa.

**Figure 1.**
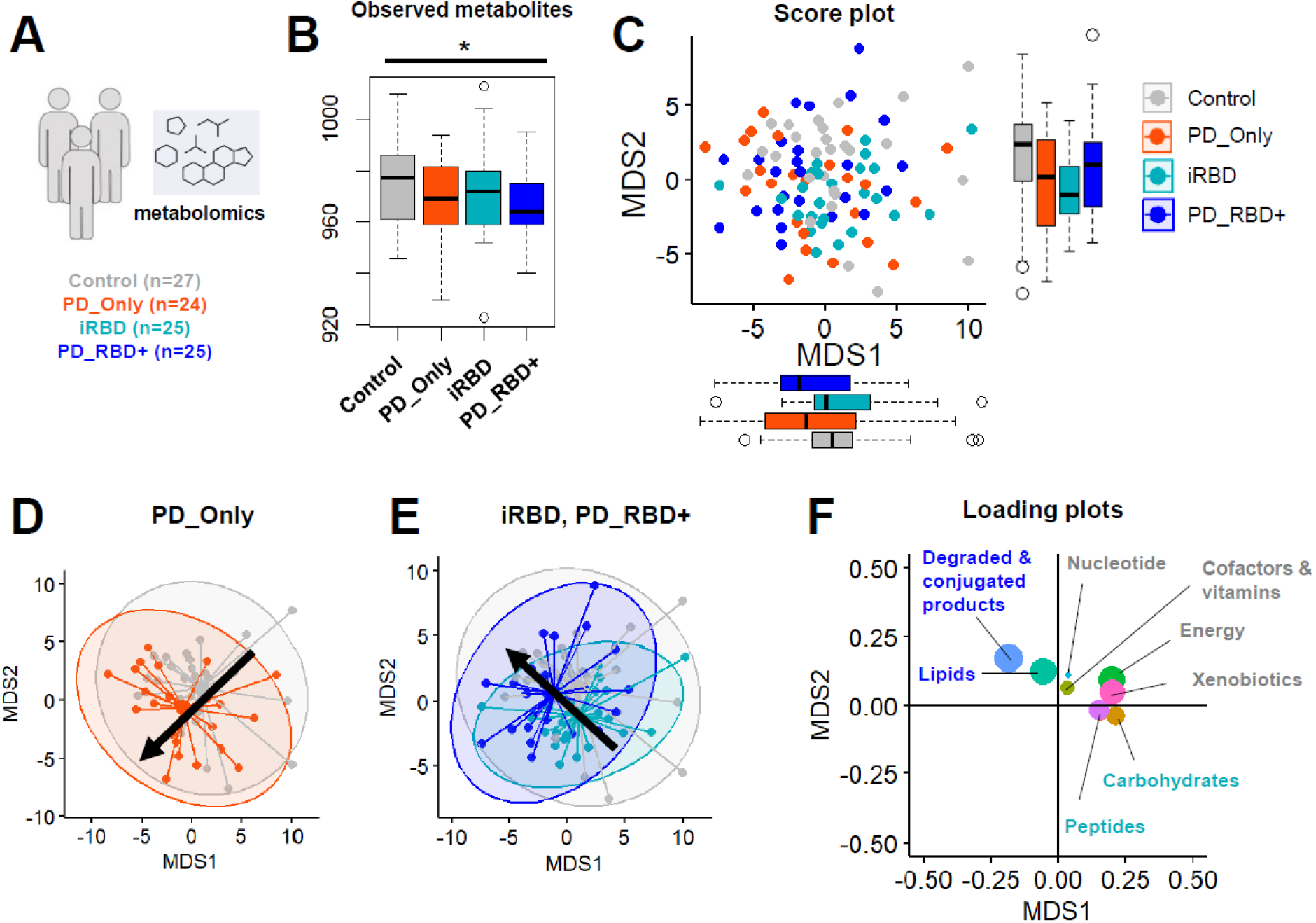
Overview of the plasma metabolic features of cohorts. **A,** Description of samples collected for untargeted metabolomics. **B,** Annotated metabolite numbers in each group from untargeted metabolomics profiles. **C,** Non-metric dimensional scaling (NMDS) ordination analysis-based plot illustrating untargeted metabolomics profiles from all groups. **D,** NMDS ordination analysis-based plot comparing untargeted metabolomics profiles between PD without RBD and controls. **E,** NMDS ordination analysis-based plot comparing untargeted metabolomics profiles between RBD groups (iRBD, PD_RBD+) and controls. **F,** NMDS loading plot summarized by pathways. Data presented as box plots show minimum, 25% quartile, median, 75% quartile, maximum, \**P* < 0.05 (Wilcoxon rank-sum tests)

Regarding metabolite analysis, a decrease in the total number of detected metabolites was observed in the diseased groups compared to healthy controls (Wilcoxon rank-sum tests p-values < 0.05) (**Figure 1B**). To gain insight into the general similarity or dissimilarity in metabolite profiles between groups, a non-metric multidimensional scaling (NMDS) plot from untargeted metabolomics analysis revealed a distinct shift in the metabolite profile in subjects with PD or RBD when compared to controls (**Figure 1C, 1D, and 1E**). We examined the loadings of the given ordination plot and summarized its loading at the pathway level (**Figure 1F**). We found that top-left directions were associated with lipids and degraded products, including bile acids and bilirubin products, which are linked to iRBD and PD_RBD+ samples in their ordination plot. Top-right directions, opposite to PD_Only samples, were associated with nucleotides, cofactors/vitamins, energy metabolites, and xenobiotics, indicating a reduction in energy metabolism and xenobiotic levels in PD_Only samples (**Figure 1F**).

### Plasma metabolic features of PD subtypes focusing on RBD

Considering that previous plasma metabolic profiling of PD patients reflects mixed profiles from PD without RBD (PD_Only) and PD with RBD (PD_RBD+) and acknowledging that RBD is a pivotal prodromal marker of PD, we have grouped PD_RBD+ and iRBD into the category “RBD group.” This grouping is undertaken to more effectively distinguish between plasma metabolic features associated with PD_Only (“brain-first PD”) and PD_RBD+ (“body-first PD”) subtypes. To identify metabolites that could predict the RBD group and PD_Only subtypes, we conducted metabolome-wide associations with clinical groups using generalized fold changes and Wilcoxon rank-sum tests (**Figure 2A, Table S1-S3**). Comparing all significantly altered metabolites (p-values < 0.05) in any of the clinical groups (PD_Only, iRBD or PD_RBD+) versus healthy controls, we observed high correlations between iRBD and PD_RBD+ groups (Pearson’s correlation coefficients = 0.510, p-value < 0.001, **Figure 2A**), indicating similar enrichments of disease metabolites in both groups. Further comparisons between all RBD groups versus control (**Figure 2B and D, Table S4**) and PD_Only group by generalized fold changes and Wilcoxon rank-sum tests (**Figure 2C and E, Table S5**) revealed RBD-specific enriched metabolites, including secondary bile acids (lithocholate sulfate, glycolithocholate), p-cresol sulfate, and phenylacetylglutamine (PAG), which are of gut microbial origin (10), Conversely, enriched metabolites in PD_Only included glucose and cortisol, while caffeine levels were decreased. Although previous studies have reported increased plasma levels of microbial p-cresol sulfate and PAG in PD (14, 16), we did not observe these in our PD_Only group but found them in the PD_RBD+ group, supporting the importance of considering PD subtypes for accurate metabolite profiling. Reductions in caffeine metabolites, including caffeine, theobromine, and 1-methylurate, were consistent features of PD, aligning with other studies (14, 26–28). We also examined enriched metabolic pathways of significantly altered metabolites in all RBD groups (**Additional Table 1**) and PD_Only group (**Additional Table 2**), identifying benzoate, sterol, histidine and tyrosine metabolites as significantly altered in the RBD group, whereas xanthine metabolites were significantly altered in PD_Only group. This differential enrichment of metabolic pathways underscores the distinct metabolic signatures between RBD groups and PD_Only group.

**Figure 2.**
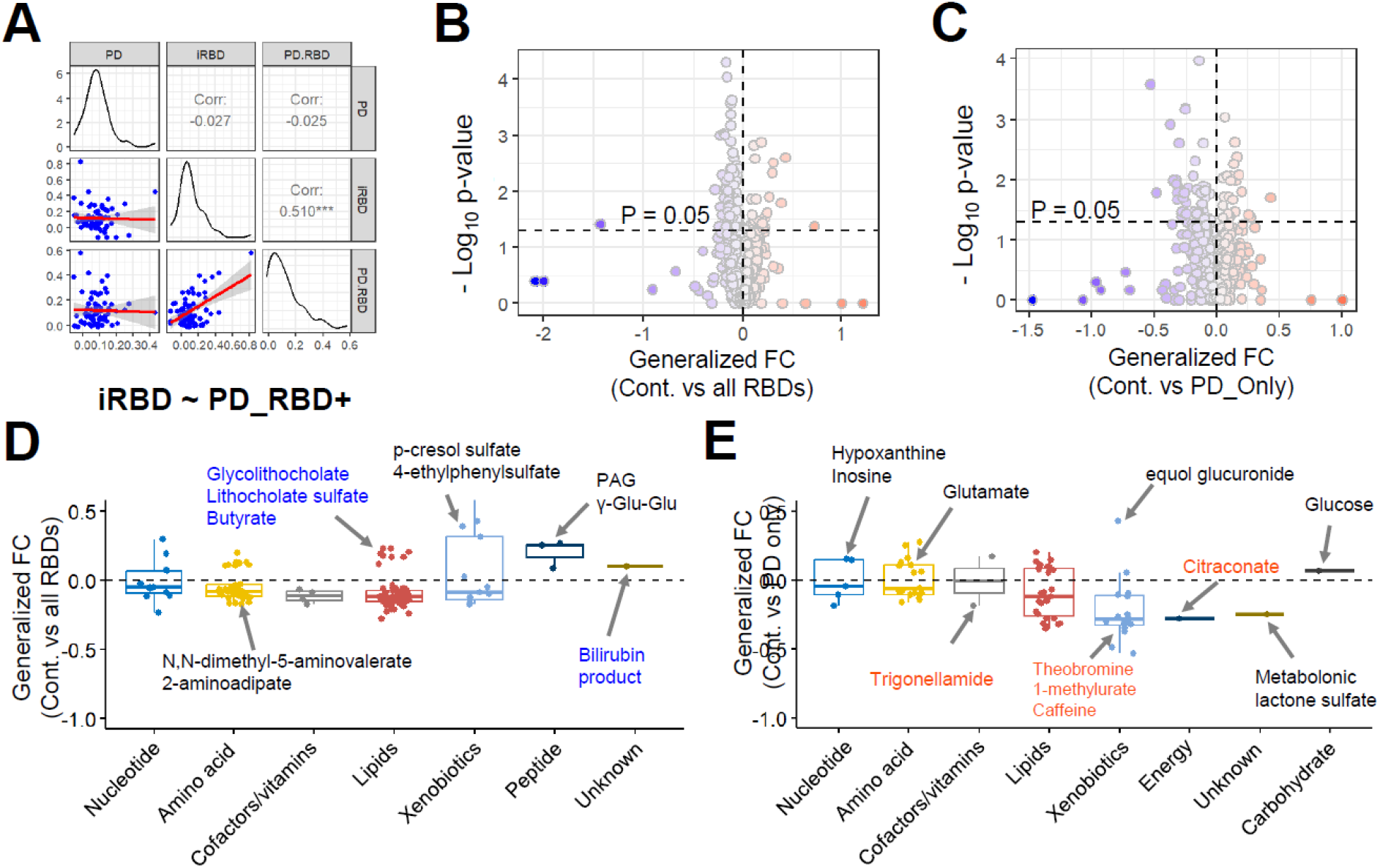
Metabolites discriminating RBD group and PD without RBD. **A,** Comparison of generalized fold changes of significantly altered metabolites (P < 0.05) from PD_Only group, iRBD group or PD_RBD+ group versus controls. Pearson’s correlation coefficients (*** *P* < 0.001) of generalized fold changes from three groups versus controls were compared. **B,** Generalized fold changes and statistical significance (-log 10 p-values) for metabolites altered between RBD groups (iRBD and PD_RBD+) vs control. **C,** Generalized fold changes and statistical significance (-log 10 p-values) for metabolites altered between PD_Only groups vs control. **D,** Generalized fold changes of metabolites significantly altered (*P* < 0.05) between RBD groups vs control summarized by pathway classes. Metabolites corresponding to degraded/conjugated products and lipids were colored blue. **E,** generalized fold changes of metabolites significantly altered (*P* < 0.05) between PD_Only vs control, by pathway classes. All statistical significance of testing significantly altered metabolites were conducted by non-parametric Wilcoxon rank-sum tests. Metabolites corresponding to xenobiotics, cofactors/vitamins and energy metabolites were colored red.

### Enrichment of feces-derived metabolites in the plasma of subjects with RBD

Given that pathologies are believed to originate from the gut in the body-first PD subtype, our investigation focused on identifying the potential origins of metabolites specific to PD_Only or RBD group. For this, we extracted PD_Only-enriched metabolites (n = 24), including cortisol, and RBD group-enriched metabolites (n = 29), including p-cresol sulfate (**Figure 3A-C and Table S6**). We then selected RBD metabolites (RM) and PD_Only metabolites (POM), which did not share each other (3 shared metabolites) for the prediction of their origins and associated bio-specimens (**Figure 3D and E** and **Table S7 and S8)** based on the Human Metabolome Database (HMDB) (21). Interestingly, among RBD group-enriched metabolites with HMDB annotations, 90% were predicted to originate from feces, while 55% were from feces in the PD_Only groups (**Figure 3D and Table S6**). This was further supported by a stronger association of microbial origin with metabolites in the RBD group compared to those in PD_Only samples (Hypergeometric test, p-value < 0.05, **Figure 3E**). One of the prominent metabolites of fecal and microbial origins, p-cresol sulfate, was elevated in iRBD and maintained its elevation in PD_RBD+ but not in PD_Only groups (**Figure 3C**), supporting PD_RBD+ as a body-first PD subtype. Moreover, when examining associations with diseases, metabolites enriched in PD_Only were associated with neuro-psychiatric diseases (i.e., schizophrenia); however, those enriched in the RBD group were associated with colon cancer and inflammatory bowel disease, both of which are closely related to the gut microbiome dysbiosis (**Figure 3G and F, Table S9 and S10**).

**Figure 3.**
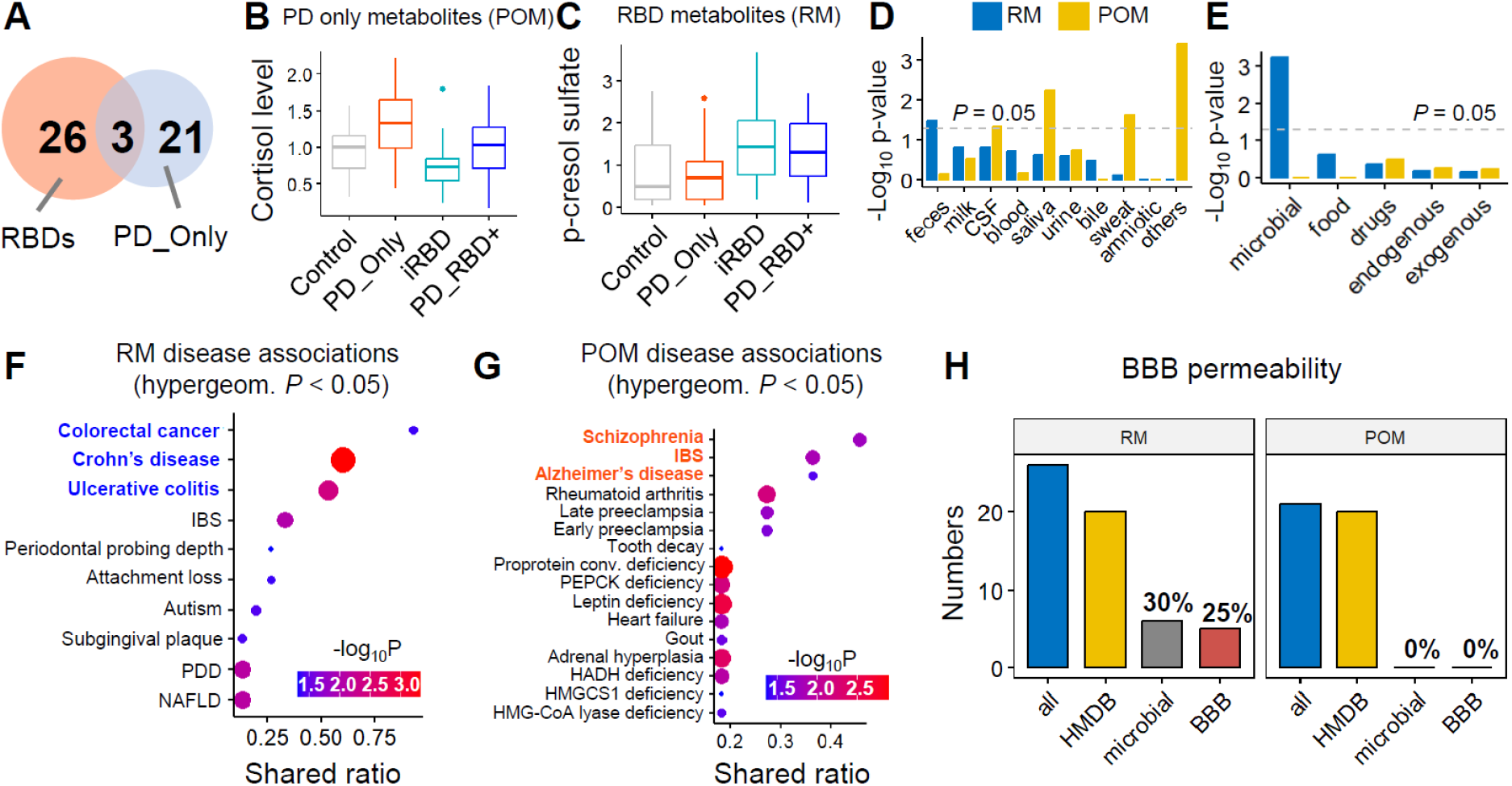
Characterizations of significantly enriched metabolites in RBD (iRBD and PD_RBD+) group and PD_Only group. **A**, Venn diagram of enriched metabolites in RBD (iRBD and PD_RBD+) group and PD_Only group. **B**, Relative abundance level of cortisol in different groups - a metabolite significantly enriched in PD_Only group (POM). **C**, Relative abundance level of p-cresol level in different groups - a metabolite significantly enriched in RBD group (RM). **D**, Significantly enriched known biospecimen type based on Human Metabolome Database (HMDB) annotations (Hypergeometric test p-values < 0.05). RBD metabolites (RM) were significantly enriched among feces metabolites, whereas PD_Only metabolites (POM) were significantly enriched among saliva, sweat, and CSF metabolites. **E**, Significantly enriched the origins of metabolites based on Human Metabolome Database (HMDB) annotations (Hypergeometric tests *P* < 0.05). RBD metabolites (RM) were significantly enriched among microbial metabolites. **F,** Significantly enriched disease metabolites of given RBD metabolites (RM) (*P* < 0.05). Statistical significance (hypergeometric tests p-values) and fraction of shared RM metabolites with disease metabolites (shared ratio) was shown by color scales and circle sizes. **G,** Significantly enriched disease metabolites of given PD_Only metabolites (POM) (*P* < 0.05). Statistical significance (hypergeometric tests p-values) and fraction of shared RM metabolites with disease metabolites (shared ratio) was shown by color scales and circle sizes. **H**, BBB permeability of RBD metabolites (RM) and PD_Only metabolites (POM). We checked metabolites of HMDB annotations from RMs and POMs and those having microbial origins (30% and 0%, respectively). For those metabolites having microbial origins, we checked BBB permeability by the *lightBBB* prediction model (25% and 0% for RM and POM, respectively).

### Prediction of metabolites of microbial origins with ability to penetrate blood-brain barrier

In the body-first PD subtype, where pathologies from the gut are suggested to invade the CNS via the vagus nerve (2), the potential of gut microbiome-derived metabolites in direct penetration to the CNS is less emphasized. We thus determined the probability of blood-brain barrier (BBB) permeability using LightBBB (22) among metabolites with microbial origins (**Table S11)**. RBD group-enriched p-cresol sulfate (**Figure 3H**) with microbial origin was predicted to penetrate the BBB, consistent with a study showing elevated p-cresol sulfate levels in the cerebrospinal fluid (CSF) of PD patients (29). Given that CSF represents the closet approximation of extracellular space of the CNS, these results suggest that RBD-enriched p-cresol sulfate as a potential mediator in the progression of PD in the body-first PD subtype.

### Establishment and evaluation of prediction models of PD clinical groups

We proceeded to assess the marker potential of discriminative metabolites identified in PD clinical groups, specifically PD_Only metabolites and RBD metabolites (**Figure 4, Table S13-S15**). Firstly, we constructed ROC curves using POM and RM metabolites to classify clinical groups versus healthy controls. Of note, among POM and RM metabolites, we excluded drug compounds, their derivatives or xenobiotics compounds from the potential markers of classification models due to their exogenous nature. Based on highest generalized fold changes, we identified three markers from POM metabolites – glucose, cortisol, and sphingomyelin (d18:1/20:2, d18:2/20:1, d16:1/22:2) - showing high AUROC values (> 0.725) to classify the PD_Only group from healthy controls (**Figure 4A, B, and C**). In addition, three markers from RM metabolites – butyrate/iso-butyrate, p-cresol sulfate, and hydrocinnamate - showed high AUROC values (> 0.706) to classify iRBD/PD_RBD+ groups from healthy controls (**Figure 4D, E, and F**). Furthermore, we included known markers from other metabolomics studies of PD, which lacked clinical clustering by RBD manifestation, such as indoleacetate, phenylacetylglutamine (PAG), and free fatty acid (10:0). Although they showed high AUROC values to classify the PD_Only group from control or iRBD/PD_RBD+ group from control, their values were still lower than those from POM and RM metabolites (**Figure 4G, H, and I**), underscoring the superior marker potential of the three metabolites we identified. Next, we established logistic regression models using PD markers – glucose, cortisol, and sphingomyelin (**Figure 4J and M**), as well as RBD markers – butyrate, p-cresol sulfate, and hydrocinnamate (**Figure 4K, N, and O**). Interestingly, based on 10-fold cross-validations with 10 repetitions, the average AUROC values were substantially high to classify the PD_Only group from control and RBD groups from control. Moreover, we assessed the prediction accuracy of our dataset aging by the logistic regression model established (confusion matrix in **Figure 4M-O**) and found that the predicted class labels closely matched with the true class labels without biases in predicting clinical groups and controls. Consequently, our models achieved high accuracy for the diagnosis of clinical groups – 80.4% accuracy to predict the PD_Only group, 69.2% accuracy to predict the PD_RBD+ group, and 74% accuracy to predict any RBD group from control.

**Figure 4.**
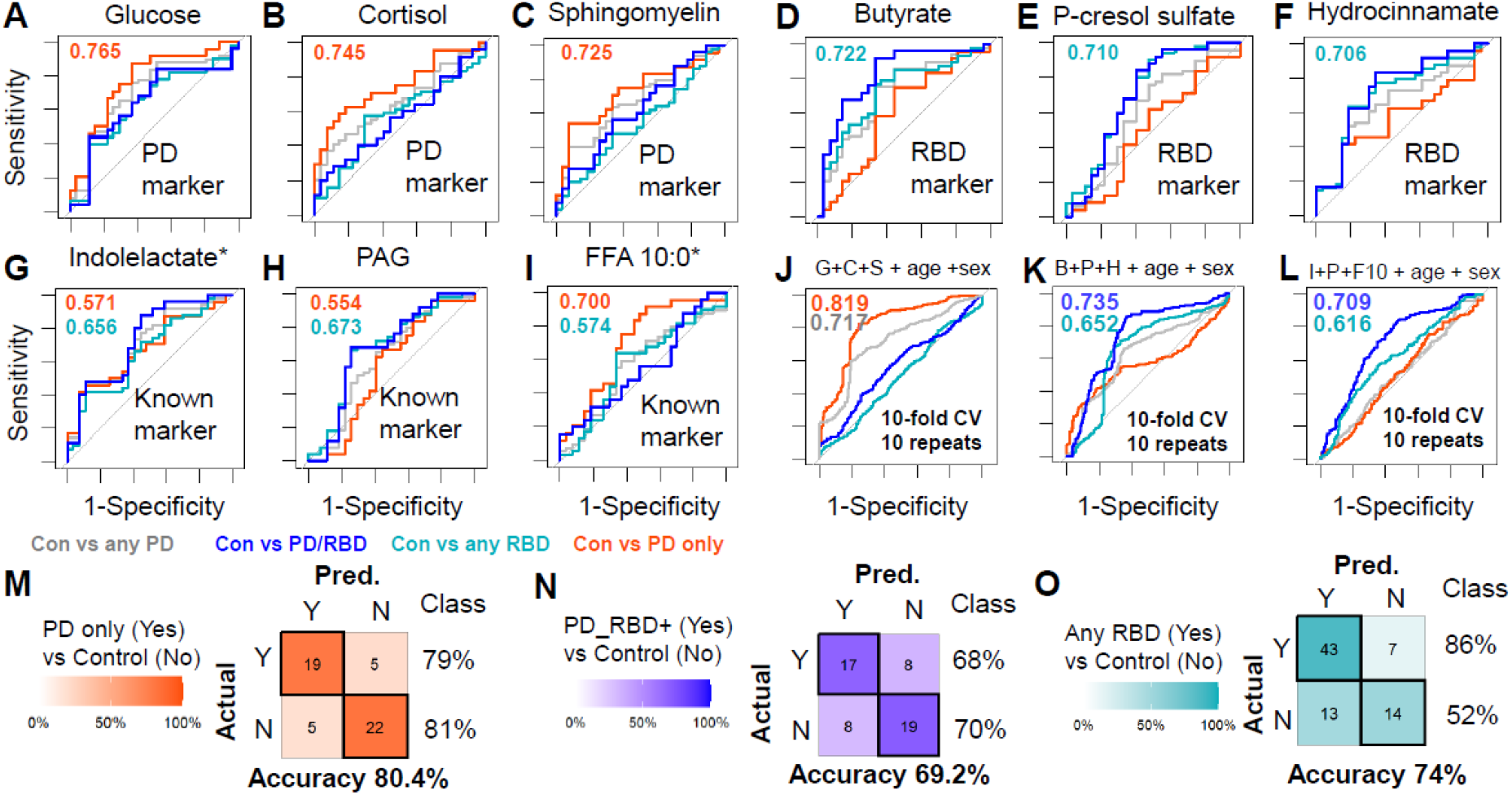
evaluation of most discriminative markers from PD_Only metabolites (POM) and RBD metabolites (RM) for the classification of clinical groups. **A-C,** most discriminative markers of PD_Only group versus controls were shown (sensitivity and specificity): (A) Glucose, (B) Cortisol, and (C) Sphingomyelin. **D-F,** most discriminative markers of RBD groups versus controls were shown (sensitivity and specificity): (D) Butyrate/iso-butyrate, (E) p-cresol sulfate, and (F) hydrocinnamate. G**-J,** Known PD metabolite markers (Yaping Shao et al., *Molecular Neurodegeneration*, 2021) were shown (sensitivity and specificity): (G) indolelactate, (H) phenylacetylglutamine (PAG), (I) free fatty acids (10:0), **J-L**, performance of logistic regression models using most discriminative markers of (J) three PD_Only metabolites (PM) - glucose, cortisol, sophingomyelin, and (K); three RBD metabolites (RM) - butyrate, p-cresol sulfate, and hydrocinnamate; and three known PD marker – indoleacetate, PAG, and FFA (10:0). **M**, predicted accuracy based on logistic regression model of (J) to classify PD_Only versus control. **N**, predicted accuracy based on logistic regression model of (K) to classify PD_RBD+ versus control. **O**, predicted accuracy based on logistic regression model of (K) to classify any RBD group versus control.

## Discussion

In this study, we investigated the plasma metabolic profiles of PD subtypes, specifically focusing on RBD as a prodromal marker (**Figure 5**). Untargeted metabolomics analysis of plasma samples from the RBD group (comprising iRBD and PD with RBD) revealed distinct metabolic features compared to the PD_Only group (PD without RBD), supporting the notion of differential pathogenesis between these subtypes. When aggregating both PD subtypes into PD, our findings aligned with previous metabolomics studies, showing altered levels of metabolites including increased microbial metabolites (i.e., secondary bile acids, p-cresol sulfate and phenylacetylglutamine), cortisol, fatty acids, branched-chain amino acids, as well as caffeine in peripheral blood (30–35). However, inconsistencies between studies (32, 36, 37) underscored the heterogeneous nature of PD pathogenesis, highlighting the need for more comprehensive subtyping. While differential pathogenesis between PD with RBD and without RBD is recognized, our study is among the first to consider the presence or absence of RBD at PD onset, which may partially explain the observed discrepancies between studies. Furthermore, our results showed that plasma metabolites in the RBD group are enriched with gut microbial origins, aligning with clinical, neuroimaging, and neuropathological evidence supporting the ‘brain-first vs body-first’ hypothesis (38, 39).

**Figure 5.**
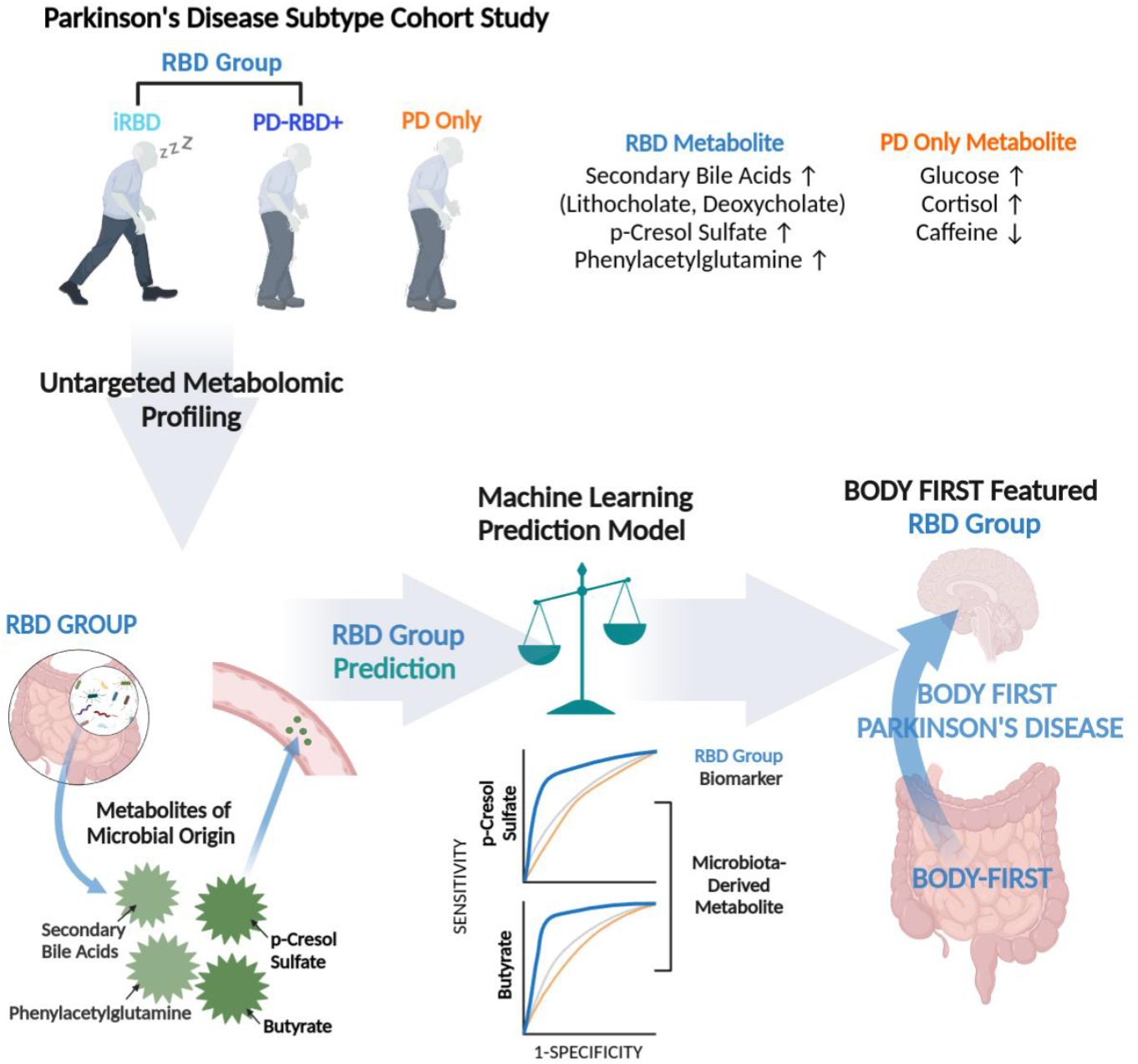
Schematic illustration of plasma metabolic profiles in PD subtypes based on RBD. This figure illustrates the plasma metabolic profiles observed in PD subtypes, categorized by the presence or absence of RBD, utilizing an untargeted metabolomics approach. Across PD cases with and without RBD, elevated levels of secondary bile acids, p-cresol sulfate, phenylacetylglutamine, and cortisol, alongside decreased caffeine, were identified, consistent with previously reported PD-associated metabolites. However, upon closer examination focusing on the presence of RBD, a significant divergence was noted. Specifically, secondary bile acids, p-cresol sulfate, and phenylacetylglutamine—metabolites originating from gut microbial activity—were distinctly evident in PD cases with RBD but absent in those without RBD. This finding suggests a potential association between PD subtypes characterized by RBD as a “body-first” type PD.

Our approach of segregating PD based on the presence or absence of RBD enabled us to infer metabolic pathways common to PD with or without RBD, specific to RBD, and specific to PD_Only, indicating both shared and distinct pathological and functional alterations. Among the common metabolic pathways identified were lysophospholipid metabolism; methionine, cysteine, SAM and Taurine metabolism; fatty acid metabolism etc. Lysophospholipids, the smallest bioactive lipids, play diverse biological roles in the brain (i.e., synapse formation, synaptic transmission, and neuron survival (40). Some lysophospholipid species are shared regardless of RBD presence, while others are distinctly enriched in PD with or without RBD (Additional Table 1 and 2). Notably, alterations in lysophosphatidylcholines have been observed in the midbrain of 6-hydroxydopamine (6-OHDA)-induced rat model of PD. Thus, the reduction in lysophospholipid species might be associated with mitochondrial impairment and neuronal degeneration in both subtypes of PD.

Overrepresented metabolic pathways in PD without RBD include metabolites associated with xanthine and purine metabolic pathways, which exhibit decreased abundance, such as caffeine, inosine, 1-methylurate, 5-acetylamino-6-formylamino-3-mehtyluracil, and 1,7-dimethylurate (Additional Table 2). It is interesting to note that dietary intake of caffeine is associated with a reduced risk for PD development in epidemiological studies (41) Lower inosine blood content in PD, as identified in our study, was also reported in the previous metabolic analysis (42). Urate, the end product of purine metabolism via inosine and xanthine, is inversely correlated with PD risk (43–46). Together with caffeine (non-selective adenosine A2A antagonists), urate has emerged as neuroprotective potential with its antioxidant capacity (47–52). PD subtyping based on the presence of RBD in our study further highlights that reduced levels of inosine, a precursor of urate, and urate itself were specific to the brain-first type of PD and were observed at the onset of the disease. However, recent phase 2 and phase 3 clinical trials of inosine in early PD (clinicaltrials.gov identifier # NCT00833690 and NCT02642393) did not yield significant differences in the rate of clinical disease progression. This underscores the importance of stratifying PD patients for this intervention, especially considering that inosine and urate levels were not affected in PD with RBD.

Metabolic pathways overrepresented in RBD groups include the long-chain polyunsaturated fatty acid metabolic pathway. Interestingly, several polyunsaturated fatty acids have shown to play an important role in the regulation of REM sleep, with omega-3 long-chain polyunsaturated fatty acid shown to improve sleep health in childhood (53). There is a significant negative correlation between sleep disturbance and the blood concentrations of long-chain polyunsaturated fatty acids (myristic, palmitic, palmitoleic, oleic, linoleic, eicosadienoic and docosahexaenoic acid) (54), with specific fatty acids such as palmitoleic and oleic acids acting as precursors for sleep-inducing compounds like oleamide and PGD2 (54). Moreover, it has been shown that treatment with DHA and EPA improves sleep disorder-related depression in PD patients (55). Thus, it is feasible to speculate that the general repression of metabolites in the long-chain polyunsaturated fatty acid metabolic pathway in RBD might reflect dysfunctional biochemical production of sleep modulators due to neurodegeneration induced by a-synuclein pathology infiltration to the lower brainstem. Distinct signatures of identified metabolites in these metabolic pathways could potentially be used to distinguish and diagnose RBD with PD related a-synuclein pathologies in the lower brainstem.

Another metabolic pathway that is overrepresented in RBD groups is benzoate metabolism. Within this pathway, metabolites such as hydrocinnamate and 4-ethylphenylsulfate, and p-cresol sulfate were substantially increased, all of which have microbial origins However, it is not clear whether these metabolites play a pathological or protective role in RBD. Hydrocinnamate, for instance, is produced by microbial metabolism of polyphenols, and serum levels of hydrocinnamate are positively associated with microbial alpha diversity, indicative of a healthy gut (56). Similarly, cinnamic acid, a related compound, has shown neuroprotective activity in a mouse model of neurodegeneration (57).

On the other hand, 4-ethylphenylsulfate (4-EPS), a microbiota-derived metabolite found at increased levels in the plasma of individuals with autism spectrum disorder, has been shown to induce anxiety-like behaviors in mice when derived from the gut and crossing the blood-brain barrier (58–60), although its impact in RBD and PD has not been investigated. p-cresol sulfate, another metabolite with a microbial origin, has been detected at higher concentrations in PD across different cohorts (14, 16). p-cresol sulfate is known to accumulate with aging and influence oxidative stress and inflammation, which may play a role in behavior disorders and neurodegeneration (61). In our study, we, for the first time, showed that p-cresol sulfate is a feature of RBD and PD with RBD but not in PD without RBD at onset. However, there might be a possibility of elevation of p-cresol sulfate in progressed PD without RBD, which needs to be investigated in the future. Given that p-cresol sulfate signatures are rather consistent in PD across different studies and its specific signals in RBD in our study, we speculated that p-cresol producing bacteria and bacterial enzymes responsible for producing p-cresol would be detected in subsets of PD patients but still showed significance among heterogeneous PD populations. We thus analyzed the abundance of p-cresol producer, *Clostridioides difficile*, and *hpdB* and *hpdC* (encoding 4-hydroxyphenylacetate decarboxylase responsible for producing p-cresol) using whole-genome shotgun sequencing datasets of fecal samples from 491 individuals with PD and 234 neurologically healthy elderly controls (7). Our analysis showed trends towards higher levels of *C. difficile*, *hpdB*, and *hpdC* in subjects with PD compared to neurologically healthy controls and confirmed that subsets of PD patients had higher abundance of *C. difficile* with *hpdB* or *hpdC* (**Additional Figure 1A-1D**). This again emphasizes the importance of PD subtyping for better targeting and understanding idiopathic PD with heterogeneity.

In summary, our study illuminates the metabolic differences between PD subtypes, particularly distinguishing between those with and without RBD at onset. Using untargeted metabolomics, we identified unique metabolic patterns associated with RBD, supporting the body-first hypothesis of PD pathogenesis and suggesting plasma metabolites, particularly of gut microbial origin, as potential biomarkers for subtype detection. Moreover, the resemblance in metabolic profiles between individuals with iRBD and those with RBD in PD hints at the potential utility of metabolic biomarkers in the early diagnosis of prodromal PD. This is significant as it presents an opportunity for the early identification of PD risk among individuals experiencing sleep disturbances, potentially enabling more effective treatment and prevention of ongoing midbrain dopaminergic neurodegeneration. While providing valuable insights, limitations such as sample size and focus on plasma metabolites require validation in larger cohorts and fecal microbiome samples. Nonetheless, our findings emphasize the need for deep phenotyping for better PD diagnosis and potentially pave the way for tailored personalized treatment based on accurate subtype diagnosis.

## Additional information

### Additional Table 1

Enriched metabolic pathways that are associated with significantly altered metabolites in the plasma of RBD patients as compared to that of age matched healthy control. Decreased metabolites in RBD patients are colored in blue, while increased metabolites in RBD patients are colored in red. # indicates metabolites that are commonly altered in both Ctrl vs. RBD and Ctrl vs PD comparison.

### Additional Table 2

Enriched metabolic pathways that are associated with significantly altered metabolites in the plasma of PD patients as compared to that of age matched healthy control. Decreased metabolites in RBD patients are colored in blue, while increased metabolites in RBD patients are colored in red. # indicates metabolites that are commonly altered in both Ctrl vs. RBD and Ctrl vs PD comparison.

**Additional Figure 1.**
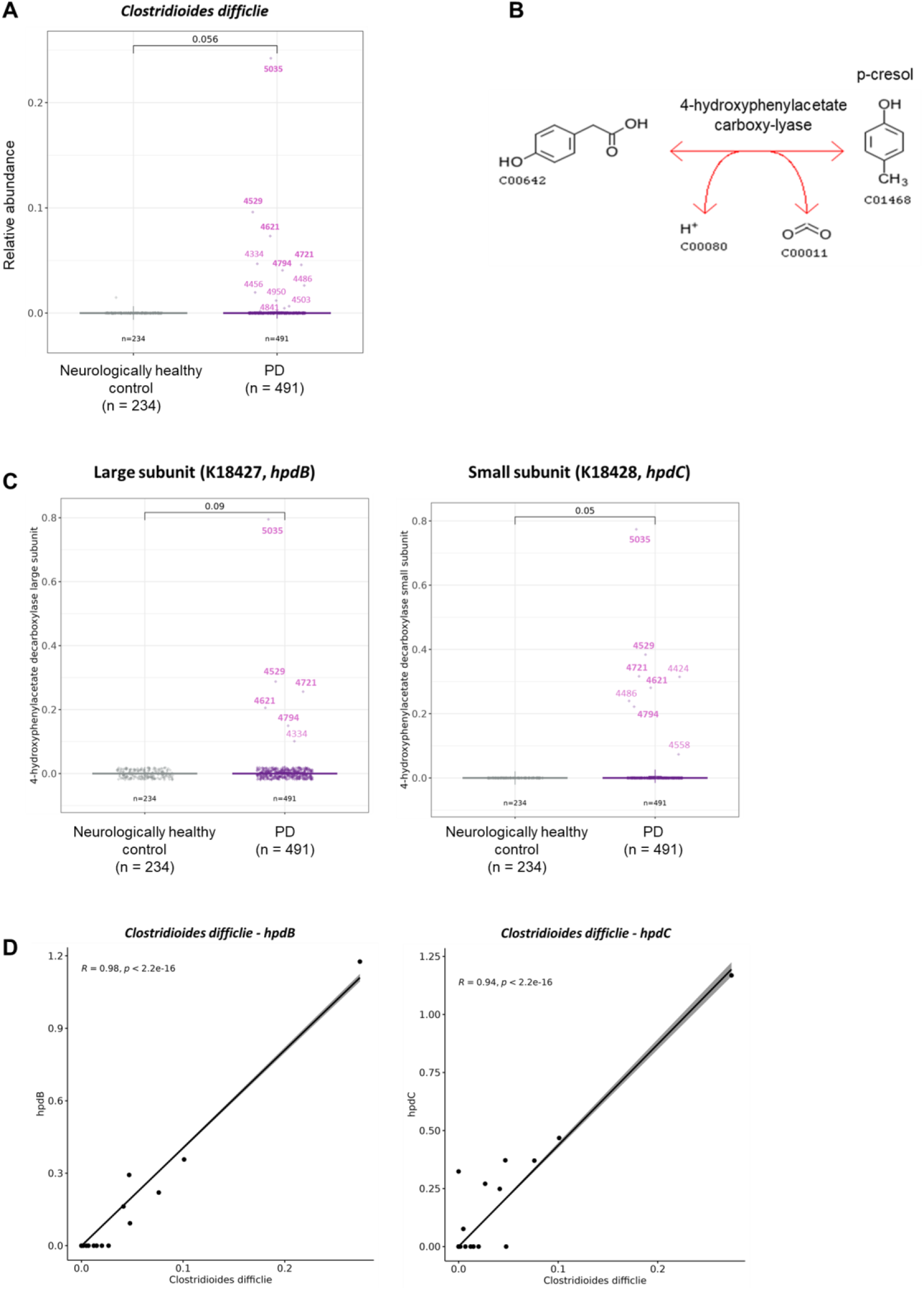
Identification of p-cresol signatures in the gut microbiome of PD. **A,** Abundance of known p-cresol producer, *Clostridioides difficile.* **B,** Microbial p-cresol production pathway via 4-hydroxyphenylacetate decarboxylase (hpd). **C,** Abundance of *hpdB* (large subunit, KEGG ortholog K18427) and *hpdC* (small subunit, KEGG ortholog K18428). *P*-values were determined by Wilcoxon rank-sum test. **D,** Pearson’s correlation between *hpdB*, *hpdC* counts and relative abundance of *C. difficile.* Each dot represents an individual data point, and numbers indicate patient ID.

## Supplementary Information

**Table S1.** Significantly altered metabolites in PD_Only group, compared to control (*P* < 0.05)

**Table S2.** Significantly altered metabolites in iRBD group, compared to control (*P* < 0.05)

**Table S3.** Significantly altered metabolites in PD_RBD+ group, compared to control (*P* < 0.05)

**Table S4.** Wilcoxon rank-sum tests and generalized fold change statistics for comparing control vs all RBD groups (iRBD & PD_RBD+)

**Table S5.** Wilcoxon rank-sum tests and generalized fold change statistics for comparing control vs PD_Only group

**Table S6.** Lists of enriched metabolites in PD_Only (POM) and RBD groups (RM) and those enriched in both PD_Only and RBD group

**Table S7.** Enrichment tests of metabolite origins for given PD_Only metabolites (POM) and RBD metabolites (RM)

**Table S8.** Enrichment tests of bio-specimen types for given PD_Only metabolites (POM) and RBD metabolites (RM)

**Table S9.** The list of BBB-permeable metabolites from RBD metabolites (RM) with microbial origins

**Table S10.** Enriched disease metabolites to PD_Only metabolites (POM)

**Table S11.** Enriched disease metabolites to RBD metabolites (RM)

**Table S13.** AUROC values of PD_Only metabolites (POM)

**Table S14.** AUROC values of RBD metabolites (RM)

**Table S15.** AUROC values of known Parkinson’s disease metabolite markers.

## Availability of data and materials

Further information and requests for resources should be directed to and will be fulfilled by the Lead Contact, HJ Kim.

## Ethics approval and consent to participate

The study protocol received approval from the Institutional Review Board of Seoul National University Hospital (IRB No. 2207-085-1340), and informed consent was waived by the board.

## References

1. Morris HR, Spillantini MG, Sue CM, Williams-Gray CH. The pathogenesis of Parkinson’s disease. Lancet. 2024;403(10423):293–304.

2. Berg D, Borghammer P, Fereshtehnejad SM, Heinzel S, Horsager J, Schaeffer E, Postuma RB. Prodromal Parkinson disease subtypes - key to understanding heterogeneity. Nature Reviews Neurology. 2021;17(6):349–61.

3. Braak H, Del Tredici K, Rüb U, de Vos RAI, Steur ENHJ, Braak E. Staging of brain pathology related to sporadic Parkinson’s disease. Neurobiology of Aging. 2003;24(2):197–211.

4. Horsager J, Andersen KB, Knudsen K, Skjærbæk C, Fedorova TD, Okkels N, et al. Brain-first versus body-first Parkinson’s disease: a multimodal imaging case-control study. Brain. 2020;143(10):3077–88.

5. Cersosimo MG, Benarroch EE. Pathological correlates of gastrointestinal dysfunction in Parkinson’s disease. Neurobiology of Disease. 2012;46(3):559–64.

6. Romano S, Savva GM, Bedarf JR, Charles IG, Hildebrand F, Narbad A. Meta-analysis of the Parkinson’s disease gut microbiome suggests alterations linked to intestinal inflammation. Npj Parkinsons Disease. 2021;7(1).

7. Wallen ZD, Demirkan A, Twa G, Cohen G, Dean MN, Standaert DG, et al. Metagenomics of Parkinson’s disease implicates the gut microbiome in multiple disease mechanisms. Nat Commun. 2022;13(1):6958.

8. Huang B, Chau SWH, Liu YP, Chan JWY, Wang J, Ma SL, et al. Gut microbiome dysbiosis across early Parkinson’s disease, REM sleep behavior disorder and their first-degree relatives. Nature Communications. 2023;14(1).

9. Cryan JF, Dinan TG. Mind-altering microorganisms: the impact of the gut microbiota on brain and behaviour. Nature Reviews Neuroscience. 2012;13(10):701–12.

10. Koh A, Backhed F. From Association to Causality: the Role of the Gut Microbiota and Its Functional Products on Host Metabolism. Mol Cell. 2020;78(4):584–96.

11. Sampson TR, Debelius JW, Thron T, Janssen S, Shastri GG, Ilhan ZE, et al. Gut Microbiota Regulate Motor Deficits and Neuroinflammation in a Model of Parkinson’s Disease. Cell. 2016;167(6):1469-+.

12. Roede JR, Uppal K, Park Y, Lee K, Tran V, Walker D, et al. Serum Metabolomics of Slow vs. Rapid Motor Progression Parkinson’s Disease: a Pilot Study. Plos One. 2013;8(10).

13. Troisi J, Landolfi A, Cavallo P, Marciano F, Barone P, Amboni M. Metabolomics in Parkinson’s disease. Advances in Clinical Chemistry, Vol 104. 2021;104:107–49.

14. Shao YP, Li TB, Liu ZY, Wang XL, Xu XJ, Li S, et al. Comprehensive metabolic profiling of Parkinson’s disease by liquid chromatography-mass spectrometry. Molecular Neurodegeneration. 2021;16(1).

15. Shao YP, Le WD. Recent advances and perspectives of metabolomics-based investigations in Parkinson’s disease. Molecular Neurodegeneration. 2019;14.

16. Paul KC, Zhang KR, Walker DI, Sinsheimer J, Yu Y, Kusters C, et al. Untargeted serum metabolomics reveals novel metabolite associations and disruptions in amino acid and lipid metabolism in Parkinson’s disease. Molecular Neurodegeneration. 2023;18(1).

17. Postuma RB, Poewe W, Litvan I, Lewis S, Lang AE, Halliday G, et al. Validation of the MDS clinical diagnostic criteria for Parkinson’s disease. Mov Disord. 2018;33(10):1601–8.

18. Postuma RB, Arnulf I, Hogl B, Iranzo A, Miyamoto T, Dauvilliers Y, et al. A single-question screen for rapid eye movement sleep behavior disorder: a multicenter validation study. Mov Disord. 2012;27(7):913–6.

19. Feng J, Meyer CA, Wang Q, Liu JS, Shirley Liu X, Zhang Y. GFOLD: a generalized fold change for ranking differentially expressed genes from RNA-seq data. Bioinformatics. 2012;28(21):2782–8.

20. Xu S, Zhan L, Tang W, Wang Q, Dai Z, Zhou L, et al. MicrobiotaProcess: A comprehensive R package for deep mining microbiome. Innovation (Camb). 2023;4(2):100388.

21. Wishart DS, Guo A, Oler E, Wang F, Anjum A, Peters H, et al. HMDB 5.0: the Human Metabolome Database for 2022. Nucleic Acids Res. 2022;50(D1):D622–D31.

22. Shaker B, Yu MS, Song JS, Ahn S, Ryu JY, Oh KS, Na D. LightBBB: computational prediction model of blood-brain-barrier penetration based on LightGBM. Bioinformatics. 2021;37(8):1135–9.

23. Wallen ZD, Demirkan A, Twa G, Cohen G, Dean MN, Standaert DG, et al. Metagenomics of Parkinson’s disease implicates the gut microbiome in multiple disease mechanisms. Nature Communications. 2022;13(1):6958.

24. Blanco-Míguez A, Beghini F, Cumbo F, McIver LJ, Thompson KN, Zolfo M, et al. Extending and improving metagenomic taxonomic profiling with uncharacterized species using MetaPhlAn 4. Nature Biotechnology. 2023;41(11):1633–44.

25. Beghini F, McIver LJ, Blanco-Míguez A, Dubois L, Asnicar F, Maharjan S, et al. Integrating taxonomic, functional, and strain-level profiling of diverse microbial communities with bioBakery 3. eLife. 2021;10:e65088.

26. Hatano T, Saiki S, Okuzumi A, Mohney RP, Hattori N. Identification of novel biomarkers for Parkinson’s disease by metabolomic technologies. Journal of Neurology Neurosurgery and Psychiatry. 2016;87(3):295–301.

27. Fujimaki M, Saiki S, Li YZ, Kaga N, Taka H, Hatano T, et al. Serum caffeine and metabolites are reliable biomarkers of early Parkinson disease. Neurology. 2018;90(5):E404-+.

28. Takeshige-Amano H, Saiki S, Fujimaki M, Ueno SI, Li YZ, Hatano T, et al. Shared Metabolic Profile of Caffeine in Parkinsonian Disorders. Movement Disorders. 2020;35(8):1438–47.

29. Sankowski B, Ksiezarczyk K, Rackowska E, Szlufik S, Koziorowski D, Giebultowicz J. Higher cerebrospinal fluid to plasma ratio of -cresol sulfate and indoxyl sulfate in patients with Parkinson’s disease. Clinica Chimica Acta. 2020;501:165–73.

30. Paul KC, Zhang K, Walker DI, Sinsheimer J, Yu Y, Kusters C, et al. Untargeted serum metabolomics reveals novel metabolite associations and disruptions in amino acid and lipid metabolism in Parkinson’s disease. Mol Neurodegener. 2023;18(1):100.

31. Breen DP, Vuono R, Nawarathna U, Fisher K, Shneerson JM, Reddy AB, Barker RA. Sleep and circadian rhythm regulation in early Parkinson disease. JAMA Neurol. 2014;71(5):589–95.

32. Chen SJ, Chen CC, Liao HY, Wu YW, Liou JM, Wu MS, et al. Alteration of Gut Microbial Metabolites in the Systemic Circulation of Patients with Parkinson’s Disease. J Parkinsons Dis. 2022;12(4):1219–30.

33. Fujimaki M, Saiki S, Li Y, Kaga N, Taka H, Hatano T, et al. Serum caffeine and metabolites are reliable biomarkers of early Parkinson disease. Neurology. 2018;90(5):e404–e11.

34. Nagesh Babu G, Gupta M, Paliwal VK, Singh S, Chatterji T, Roy R. Serum metabolomics study in a group of Parkinson’s disease patients from northern India. Clin Chim Acta. 2018;480:214–9.

35. Shao Y, Li T, Liu Z, Wang X, Xu X, Li S, et al. Comprehensive metabolic profiling of Parkinson’s disease by liquid chromatography-mass spectrometry. Mol Neurodegener. 2021;16(1):4.

36. Haglin L, Backman L. Covariation between plasma phosphate and daytime cortisol in early Parkinson’s disease. Brain Behav. 2016;6(12):e00556.

37. Shin C, Lim Y, Lim H, Ahn TB. Plasma Short-Chain Fatty Acids in Patients With Parkinson’s Disease. Mov Disord. 2020;35(6):1021–7.

38. Borghammer P, Horsager J, Andersen K, Van Den Berge N, Raunio A, Murayama S, et al. Neuropathological evidence of body-first vs. brain-first Lewy body disease. Neurobiol Dis. 2021;161:105557.

39. Horsager J, Andersen KB, Knudsen K, Skjaerbaek C, Fedorova TD, Okkels N, et al. Brain-first versus body-first Parkinson’s disease: a multimodal imaging case-control study. Brain. 2020;143(10):3077–88.

40. Fukushima N, Weiner JA, Chun J. Lysophosphatidic acid (LPA) is a novel extracellular regulator of cortical neuroblast morphology. Dev Biol. 2000;228(1):6–18.

41. Zhao Y, Lai Y, Konijnenberg H, Huerta JM, Vinagre-Aragon A, Sabin JA, et al. Association of Coffee Consumption and Prediagnostic Caffeine Metabolites With Incident Parkinson Disease in a Population-Based Cohort. Neurology. 2024;102(8):e209201.

42. LeWitt PA, Li J, Lu M, Guo L, Auinger P, Parkinson Study Group DI. Metabolomic biomarkers as strong correlates of Parkinson disease progression. Neurology. 2017;88(9):862–9.

43. Annanmaki T, Muuronen A, Murros K. Low plasma uric acid level in Parkinson’s disease. Mov Disord. 2007;22(8):1133–7.

44. Andreadou E, Nikolaou C, Gournaras F, Rentzos M, Boufidou F, Tsoutsou A, et al. Serum uric acid levels in patients with Parkinson’s disease: Their relationship to treatment and disease duration. Clinical Neurology and Neurosurgery. 2009;111(9):724–8.

45. Davis JW, Grandinetti A, Waslien CI, Ross GW, White LR, Morens DM. Observations on serum uric acid levels and the risk of idiopathic Parkinson’s disease. American Journal of Epidemiology. 1996;144(5):480–4.

46. Weisskopf MG, O’Reilly E, Chen H, Schwarzschild MA, Ascherio A. Plasma urate and risk of Parkinson’s disease. American Journal of Epidemiology. 2007;166(5):561–7.

47. Ames BN, Cathcart R, Schwiers E, Hochstein P. Uric acid provides an antioxidant defense in humans against oxidant- and radical-caused aging and cancer: a hypothesis. Proc Natl Acad Sci U S A. 1981;78(11):6858–62.

48. Yeum KJ, Russell RM, Krinsky NI, Aldini G. Biomarkers of antioxidant capacity in the hydrophilic and lipophilic compartments of human plasma. Archives of Biochemistry and Biophysics. 2004;430(1):97–103.

49. Scott GS, Spitsin SV, Kean RB, Mikheeva T, Koprowski H, Hooper DC. Therapeutic intervention in experimental allergic encephalomyelitis by administration of uric acid precursors. Proceedings of the National Academy of Sciences of the United States of America. 2002;99(25):16303–8.

50. Yu ZF, Bruce-Keller AJ, Goodman Y, Mattson MP. Uric acid protects neurons against excitotoxic and metabolic insults in cell culture, and against focal ischemic brain injury in vivo. Journal of Neuroscience Research. 1998;53(5):613–25.

51. Romanos E, Planas AM, Amaro S, Chamorro A. Uric acid reduces brain damage and improves the benefits of rt-PA in a rat model of thromboembolic stroke. Journal of Cerebral Blood Flow and Metabolism. 2007;27(1):14–20.

52. Haberman F, Tang SC, Arumugam TV, Hyun DH, Yu QS, Cutler RG, et al. Soluble neuroprotective antioxidant uric acid analogs ameliorate ischemic brain injury in mice. Neuromolecular Medicine. 2007;9(4):315–23.

53. Dai Y, Liu J. Omega-3 long-chain polyunsaturated fatty acid and sleep: a systematic review and meta-analysis of randomized controlled trials and longitudinal studies. Nutr Rev. 2021;79(8):847–68.

54. Irmisch G, Schlafke D, Gierow W, Herpertz S, Richter J. Fatty acids and sleep in depressed inpatients. Prostaglandins Leukot Essent Fatty Acids. 2007;76(1):1–7.

55. Avallone R, Vitale G, Bertolotti M. Omega-3 Fatty Acids and Neurodegenerative Diseases: New Evidence in Clinical Trials. Int J Mol Sci. 2019;20(17).

56. Wilmanski T, Rappaport N, Earls JC, Magis AT, Manor O, Lovejoy J, et al. Blood metabolome predicts gut microbiome α-diversity in humans. Nature Biotechnology. 2019;37(10):1217-+.

57. Prorok T, Jana M, Patel D, Pahan K. Cinnamic Acid Protects the Nigrostriatum in a Mouse Model of Parkinson’s Disease via Peroxisome Proliferator-Activated Receptoralpha. Neurochem Res. 2019;44(4):751–62.

58. Hsiao EY, McBride SW, Hsien S, Sharon G, Hyde ER, McCue T, et al. Microbiota Modulate Behavioral and Physiological Abnormalities Associated with Neurodevelopmental Disorders. Cell. 2013;155(7):1451–63.

59. Needham BD, Funabashi M, Adame MD, Wang Z, Boktor JC, Haney J, et al. A gut-derived metabolite alters brain activity and anxiety behaviour in mice. Nature. 2022;602(7898):647-+.

60. Needham BD, Adame MD, Serena G, Rose DR, Preston GM, Conrad MC, et al. Plasma and Fecal Metabolite Profiles in Autism Spectrum Disorder. Biological Psychiatry. 2021;89(5):451–62.

61. Sun CY, Li JR, Wang YY, Lin SY, Ou YC, Lin CJ, et al. p-Cresol Sulfate Caused Behavior Disorders and Neurodegeneration in Mice with Unilateral Nephrectomy Involving Oxidative Stress and Neuroinflammation. International Journal of Molecular Sciences. 2020;21(18).

