## Additional Tables for "Untargeted Plasma Metabolomics Unveils Distinct Metabolite Profiles in Parkinson’s Disease Subtypes: A Focus on idiopathic REM Sleep Behavior Disorders"

| **Pathways (Ctrl vs RBD)** | **# of DEMs** | **Metabolites** |
| --- | --- | --- |
| Lysophospholipid | 12 | #1-palmitoleoyl-GPI (16:1)*, #1-dihomo-linolenoyl-GPC (20:3n3 or 6)*, #1-oleoyl-GPI (18:1), #1-arachidonoyl-GPI (20:4)*, 1-linoleoyl-GPI (18:2)*, 1-palmitoyl-GPI (16:0), 1-palmitoleoyl-GPC (16:1)*, 1-oleoyl-GPC (18:1), 1-docosahexaenoyl-GPE (22:6)*, 1-linoleoyl-GPC (18:2), 1-stearoyl-GPI (18:0), 1-palmitoyl-GPC (16:0) |
| Long Chain Polyunsaturated Fatty Acid (n3 and n6) | 8 | heneicosapentaenoate (21:5n3), docosapentaenoate (n6 DPA; 22:5n6), linoleate (18:2n6), docosahexaenoate (DHA; 22:6n3), docosapentaenoate (n3 DPA; 22:5n3), docosadienoate (22:2n6)  dihomo-linolenate (20:3n3 or n6), arachidonate (20:4n6) |
| Chemical | 5 | ethyl glucuronide, iminodiacetate (IDA), perfluorohexanesulfonate (PFHxS), O-sulfo-tyrosine, sulfate* |
| Methionine, Cysteine, SAM and Taurine Metabolism | 5 | #hypotaurine, N-acetyltaurine, alpha-ketobutyrate, cystine, cystathionine |
| Benzoate Metabolism | 5 | 4-vinylphenol sulfate, benzoate, p-cresol sulfate, 4-ethylphenylsulfate, 3-phenylpropionate (hydrocinnamate) |
| Lysine Metabolism | 4 | N2-acetyl,N6-methyllysine, N-acetyl-2-aminoadipate, 2-aminoadipate, N,N-dimethyl-5-aminovalerate |
| Fatty Acid, Monohydroxy | 4 | 3-hydroxyoleate*, 2-hydroxynervonate*, 2-hydroxybehenate, 2-hydroxypalmitate |
| Histidine Metabolism | 4 | #N-acetyl-1-methylhistidine*, #imidazole lactate, N-acetylhistidine, hydantoin-5-propionate |
| Sterol | 4 | #cholesterol sulfate, 7alpha-hydroxy-3-oxo-4-cholestenoate (7-Hoca), 3beta-hydroxy-5-cholestenoate, cholesterol |
| Tyrosine Metabolism | 4 | #N-acetyltyrosine, N-formylphenylalanine, vanillylmandelate (VMA), thyroxine |

**Additional table 1.**

Enriched metabolic pathways that are associated with significantly altered metabolites in the plasma of RBD patients as compared to that of age matched healthy control. Decreased metabolites in RBD patients are colored in blue, while increased metabolites in RBD patients are colored in red. # indicates metabolites that are commonly altered in both Ctrl vs. RBD and Ctrl vs PD comparison.

| **Pathways (Ctrl vs PD)** | **# of DEMs** | **Metabolites** |
| --- | --- | --- |
| Xanthine Metabolism | 7 | theophylline, 1-methylurate, 5-acetylamino-6-formylamino-3-methyluracil, caffeine, 5-acetylamino-6-amino-3-methyluracil, 1,7-dimethylurate, theobromine |
| Fatty Acid Metabolism (Acyl Choline) | 7 | linoleoylcholine*, dihomo-linolenoyl-choline, palmitoylcholine, stearoylcholine*, oleoylcholine, docosahexaenoylcholine, arachidonoylcholine |
| Food Component/Plant | 6 | #3-formylindole, quinate, sulfate of piperine metabolite C16H19NO3 (2)*, glucuronide of piperine metabolite C17H21NO3 (4)*, sulfate of piperine metabolite C16H19NO3 (3)*  equol glucuronide |
| Lysophospholipid | 5 | #1-palmitoleoyl-GPI (16:1)*, #1-dihomo-linolenoyl-GPC (20:3n3 or 6)*, #1-oleoyl-GPI (18:1), #1-arachidonoyl-GPI (20:4)*, 1-adrenoyl-GPC (22:4)* |
| Methionine, Cysteine, SAM and Taurine Metabolism | 5 | #Hypotaurine, cysteine, S-methylcysteine, S-methylcysteine sulfoxide, S-methylmethionine |

**Additional table 2.**

Enriched metabolic pathways that are associated with significantly altered metabolites in the plasma of PD patients as compared to that of age matched healthy control. Decreased metabolites in RBD patients are colored in blue, while increased metabolites in RBD patients are colored in red. # indicates metabolites that are commonly altered in both Ctrl vs. RBD and Ctrl vs PD comparison.
